# Consistency of affective responses to naturalistic stimuli across individuals using intersubject correlation analysis based on neuroimaging data

**DOI:** 10.1101/2024.06.12.598753

**Authors:** Junhyeok Jang, Jongwan Kim

## Abstract

In this study, we used functional magnetic resonance imaging (fMRI) data obtained for naturalistic emotional stimuli to examine the consistency of neural responses among participants in specific regions related to valence. We reanalyzed fMRI data from 17 participants as they watched episodes of “Sherlock” and used emotional ratings from 125 participants. To determine regions where neural response patterns were synchronized across participants based on the pattern of valence changes, intersubject correlation analysis was conducted. As a validation analysis, multidimensional scaling was conducted to investigate emotional representation for significant regions of interest. The results revealed that the ventromedial prefrontal cortex, bilateral superior frontal cortices, left posterior cingulate cortex, thalamus, right anterior cingulate cortex, and bilateral inferior frontal cortices showed increased neural synchrony as positive scenes were presented. Also, the bilateral superior temporal gyrus and bilateral medial temporal gyrus exhibited increased neural synchrony as negative scenes were presented. Moreover, the left inferior frontal cortex and right superior frontal gyrus were found to be engaged in emotion representation and display increased neural synchrony. These findings provide insights into the differential neural responses to emotionally evocative naturalistic stimuli as compared to conventional experimental stimuli. Also, this study highlights the future potential for using intersubject correlation analysis for examining consistency of neural responses to naturalistic stimuli.

## 1. Introduction

Humans can evaluate the external world and discern between positive and negative experiences. Emotional judgments about the external world are universally important Barrett et al. (2004). Valence which represents an emotion’s subjective positive or negative quality, influences our affective responses and perspectives. Functional magnetic resonance imaging (fMRI) offers insight into neural basis of valence by examining brain activity during emotional processing. Various brain regions implicated in valence include the amygdala (Barad et al., 2006; Shin et al., 2005), medial prefrontal cortex (Baucom et al., 2012; Kim et al., 2017; Kim et al., 2020; Kringelbach, 2005; Kringelbach et al., 2003; Mesulam, 2000; Miyake et al., 2000; Wang et al., 2010), and insula (Berntson et al., 2011; Kayyal et al., 2019; Miskovic & Anderson, 2018). However, there are limitations to how precisely these conventional approaches can identify these regions in naturalistic stimuli conditions. Thus, this study aims to enhance understanding of valence processing by analyzing fMRI data from naturalistic stimuli, aiming to uncover intricate neural synchrony mechanisms in emotional processing.

### 1.1. Naturalistic stimuli

Previous neuroscientific studies of affective processing utilized controlled stimuli such as visual, auditory, and linguistic stimuli. However, concerns about ecological validity arose due to the isolated presentation of these stimuli, creating a mismatch in extrapolating experimental data to real-life reactions (Schmuckler, 2001).

To address these concerns, previous studies (Chang et al., 2021; Finn et al., 2020; Hasson et al., 2010; Sonkusare et al., 2019; van Baar et al., 2019) employed naturalistic stimuli to explore participants’ neural responses. Naturalistic stimuli are not explicitly aimed at completing a task; they are presented with dynamic, fluid transitions over time and incorporate a variety of factors such as speech, music, motion, and the context. As naturalistic stimuli are similar to perceptual experiences faced in everyday life, they possess a high degree of ecological validity. Previous fMRI studies have shown that naturalistic stimuli activate brain regions, including the ventromedial prefrontal cortex (vmPFC) (Baldassano et al., 2017; Kim et al., 2016), superior temporal sulcus (STS) (Hasson et al., 2008; Honey et al., 2012; Kim et al., 2016; Lerner et al., 2011), and visual cortex (Baldassano et al., 2017; Hasson et al., 2008; Honey et al., 2012). These regions are known to be engaged in multisensory integration and time-sensitive information processing (Hasson et al., 2008; Lerner et al., 2011).

However, naturalistic stimuli differ from experimental stimuli in their limited capacity to evoke extreme emotional changes, posing challenges for conventional fMRI analysis over prolonged durations (Hasson et al., 2010; Najafi et al., 2017; Nastase et al., 2019; Simony et al., 2016). Thus, it is necessary to develop appropriate analysis that is both sensitive and suitable for naturalistic stimuli.

### 1.2. Intersubject correlation (ISC)

Friston et al. (1995) introduced a technique to identify neural activation linked to psychological changes through a time series general linear model. This model estimates regression coefficients for the fMRI signal, and the significance of each regression equation is assessed using t-tests or F-tests to identify conditions significantly affecting fMRI signals. However, these methods face limitations due to the characteristics of fMRI data. Formulating regression equations with fMRI data could lead to issues of overfitting and multiple comparisons (Bennett et al., 2009; Nichols & Hayasaka, 2003; Nichols, 2012; Nichols & Holmes, 2002). When the number of regression variables exceeds the available data, using multiple regression coefficients may overestimate the quirks and random noise present in a specific sample, which may yield results that do not reflect the overall population. Moreover, performing statistical tests across numerous voxels increases the risk of Type I errors. While employing statistical corrections can maintain a global error probability of 0.05, excessive strictness in criteria might result in the failure to accurately identify truly significant voxels (Nichols & Hayasaka, 2003).

To address these issues, Hasson et al. (2004) developed an intersubject correlation (ISC) analysis. The ISC measures the level of neural synchrony of the regions of interest (ROI) between participants, quantified by the correlation coefficient. The fundamental assumption of this analysis is that if a specific stimulus elicits the intended behavioral or psychological changes, synchronous changes in neural activation patterns will be observed across participants in relevant brain regions (Hasson et al., 2010; Najafi et al., 2017; Nastase et al., 2019; Simony et al., 2016). The ISC can capture complex patterns of neural activation over extended periods of time by utilizing the similarity of interparticipant data (Kauppi et al., 2010). Moreover, ISC addresses overfitting issues associated with general linear models and multiple comparison problems by using the intrinsic characteristics of the data to minimize prior assumptions and hypotheses (Hejnar et al., 2007; Turner et al., 2017; Vicente et al., 2011).

Several previous studies that focus on interparticipant neural synchrony for emotions have provided insights and helped facilitate understanding of interparticipant neural synchrony (Hasson et al., 2004; Li et al., 2021; Nummenmaa et al., 2012). Despite their contributions, these studies have certain limitations. In an investigation of the synchrony of neurologic responses associated with visual perception, Hasson et al. (2004) used movie clips to examine the impact of naturalistic stimulation on neural responses. Their findings revealed robust inter-subject correlations in the visual and auditory cortices, as well as regions processing sensory information. Notably, they observed significant neural synchrony among participants in areas unrelated to sensory perception when the movie clips elicited emotional responses. They speculated that the contextual richness of naturalistic stimuli led to neural synchrony in the regions responsible for cognitive and affective functions. However, the study by Hasson et al. (2004) exclusively focused on visual information processing under naturalistic stimulation, which leaves the possibility that neural synchronization induced by sensory perception may differ from neural synchronization induced by emotional responses or cognitive function.

Nummenmaa et al. (2012) found that transitions from positive to negative movie content increased neural synchrony among participants in higher-order cognitive and affective cortexes. These regions included the precuneus, orbitofrontal cortex (OFC), insula, dorsomedial prefrontal cortex (dmPFC), visual cortex, ventral striatum, and amygdala. Conversely, increased arousal correlated with heightened neural synchrony in the intraparietal sulcus (IPS). The results indicated that the arousal network and dorsal attention network exhibited heightened neural synchronization with increased arousal, while the default mode network showed increased neural synchronization with escalating negative emotion. The study supported the hypothesis that networks involved in valence and arousal processing are distinct and dissociated. However, the stimuli presented in the experiment by Nummenmaa et al. (2012) were short in duration (1.5 to 2 minutes) and participants received a script beforehand, potentially limiting their classification as purely naturalistic stimuli.

We aim to explore the synchronization observed in neural responses within specific valence-associated regions. To achieve this goal, we re-analyzed fMRI data (Chen et al., 2017) collected while participants viewed various extended, naturalistic emotional stimuli, addressing limitations observed in the prior studies. Unlike studies that constrained participants’ responses with pre-scripted scenarios, our approach involved exposing participants to the entire first episode of the television series ‘Sherlock’ to assess fluctuations in brain activity throughout the viewing period. This allowed us to investigate natural emotional responses and their corresponding neural activations without the constraints of pre-scripted stimuli. Furthermore, we employed behavioral emotion ratings (Kim et al., 2020) measured at a same scanning time rate as the fMRI data. Our purpose was to identify areas where neural synchronization either increased or decreased in response to emotional changes using correlation coefficients between ISC values and valence ratings. Under the assumption that this approach would facilitate the identification of neural regions where participants consistently demonstrate synchronization in response to variations in emotions, our primary aim was to pinpoint brain regions closely aligned with the valence level in neural synchrony.

## 2. Methods

### 2.1. Research Data

This study reanalyzed data obtained from previous studies conducted by Chen et al. (2017) and Kim et al. (2020). The initial dataset comprised 17 participants who watched the inaugural episode of the BBC drama series “Sherlock” while undergoing MRI scanning, as outlined by Chen et al. (2017). The participants had never seen the “Sherlock” series before the study and were scanned during two sessions, each lasting 23 and 25 minutes. The functional images were acquired using a 3T Siemens Skyra scanner with a 20-channel head coil, employing a T2-weighted echo-planar imaging (EPI) pulse sequence (TR=1,500ms, TE=28ms, flip angle=64, whole-brain coverage=27 slices of 4 mm thickness, in-plane resolution=3×3, FOV=192, ascending interleaved acquisition). High-resolution whole-brain anatomical images were acquired using a standard T1-weighted 3D MPRAGE protocol.

The fMRI data of 17 participants were preprocessed using slice timing difference correction, motion correction, linear detrending, and high-pass filtering by Chen et al. (2017). Subsequently, the images were coregistered and affine transformed to the Montreal Neurological Institute 152 (MNI 152) after resampling, resulting in functional images with isotropic voxels of 3 mm. The preprocessed data were standardized, simultaneously measured for all voxels, and smoothed using a Gaussian function with a Full Half Maximum of 6 mm (Figure 1 A).

**Figure 1.**
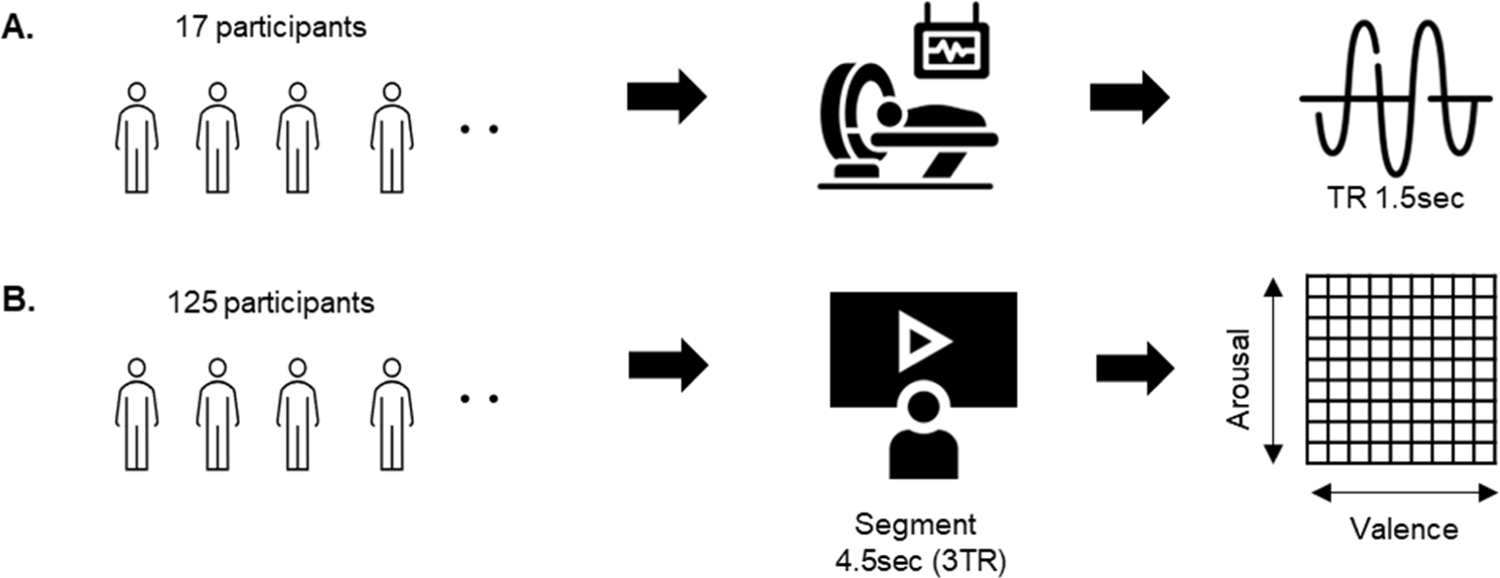
Schematic illustration of the study procedure. A. fMRI data from 17 participants B. emotion ratings data from 125 participants.

Kim et al. (2020) further processed divided the fMRI data by segmenting it into 621 segments of 4.5 seconds (3 TR) each. This segmentation strategy, aligned with multiples of the TR, aimed to synchronize the timing of behavioral ratings with the fMRI data, thereby facilitating a more comprehensive analysis. By utilizing this segment duration, they ensured ample trials while allowing participants sufficient time to thoroughly comprehend each clip and provide meaningful ratings. To account for the hemodynamic response, the fMRI data were temporally offset by 4.5 seconds from the stimulus onset, following the methodology established by Chen et al. (2017),before being averaged for each segment (Figure 1 B).

### 2.2. Behavioral emotion ratings

Kim et al. (2020) collected behavioral emotion ratings from 125 undergraduate students at the University of South Carolina (34 males, 91 females, mean age: 20.38 years) concerning the “Sherlock” episode. The assessment utilized a 9×9 emotion grid (Russell et al., 1989) to measure emotions at 4.5-second intervals, resulting in 81 emotion grids representing affective valence and arousal dimensions. In this grid, the horizontal axis represented how positive or negative the stimulus was, with a rightward rating indicating a more positive evaluation and a leftward rating indicating a more negative evaluation. As shown in Figure 1b, the vertical axis indicates the degree of arousal, where higher ratings denoted increased activation and lower ratings indicated decreased activation. For the purposes of our study, we focused on 621 segments of valence data, excluding arousal ratings, to identify neural responsiveness regions in response to changes in valence.

### 2.3. Regions of interest (ROIs)

In this study, the Automated Anatomical Labeling Atlas 3(AAL3) (Rolls et al., 2020) was used to define 170 anatomically divided regions of interest (ROIs). The AAL3 utilized voxels with a size of 2×2×2, which did not match the voxel size from the study by Chen et al. (2017). To address this, voxel reslicing was executed using the dimension coregistration function in the Statistical Parametric Mapping 12 (SPM12) toolbox, ensuring alignment with the fMRI data voxel size. The reslicing, based on categorical information within the ROIs, employed nearest neighbor interpolation. We generated 140 ROI templates using AAL 3 for analysis, excluding the cerebellum (ROIs 95-120), regions lacking white matter (ROIs 167 and 168, locus coeruleus), and ROIs 169 and 170 (raphe nucleus).

### 2.4. Intersubject correlation (ISC)

We utilized ISC to identify similar patterns of neural activation among participants in response to emotional changes (Hasson et al., 2010; Najafi et al., 2017; Nastase et al., 2019; Simony et al., 2016).

Initially, the data regarding the 17 participants were segregated into one participant S_i_ (the target participant) and the remaining participants S_n-1_ (leave-one-subject-out) (Figure 2 A). Subsequently, we extracted the voxel activation levels corresponding to a particular ROI. The voxel activation levels during each segment contained three-dimensional spatial information (x, y, z). We transformed this three-dimensional data into a one-dimensional array and organized the S_i_ data in a matrix format representing voxel activation levels for the k_th_ ROI during each segment. In the same format as S_i_ data, all S_n-1_ data matrices were averaged across participants (Figure 2 B). The next step involved examining neural synchrony for each ROI. We rearranged the segment order from negative to positive emotions based on behavioral ratings and correlated voxel activation patterns between target participant and the remaining participants for each segment (Figure 2 C). An ISC value approaching 1 suggests high synchrony, indicating regions with consistent processing across individuals, while values near 0 indicate low synchrony, reflecting regions with idiosyncratic processing or minimal encoding of stimulus information.(Nastase et al., 2019). The calculation of ISC was repeated for all segments, resulting in 621 ISC values. The entire process was iterated for all participants, producing 621 ISC values for the 17 participants (Figure 2 D). Lastly, ISC values were averaged across participants and a correlation coefficient was calculated between ISC and valence ratings for each segment. This analysis revealed that positive correlations indicated an increase in participants’ neural synchrony as stimuli became more positive. Conversely negative correlations indicated an increase in participants’ neural synchrony as the stimuli became more negative. This process was repeated for all 140 ROIs, yielding correlation coefficients between ISC and valence ratings for each ROI (Figure 2 E).

**Figure 2.**
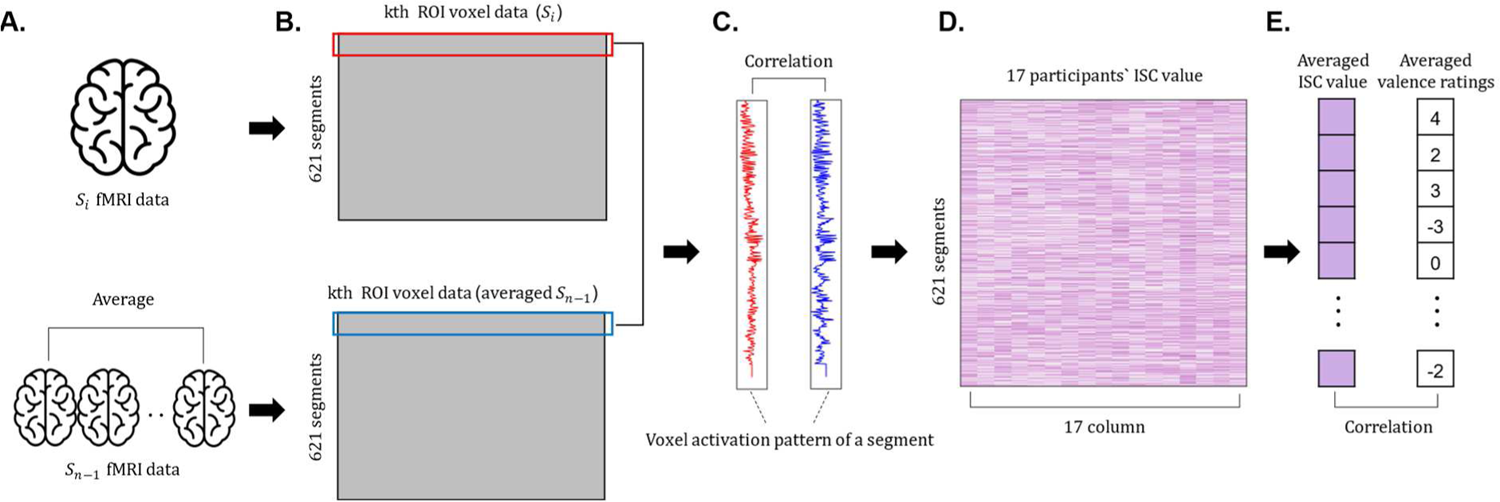
Illustration of the procedure for calculating the ISC-valence ratings correlation coefficient for ROIs

To evaluate the significance of the correlation coefficient between ISC and valence ratings for a specific ROI, we performed a permutation test. The permutation test constructs an empirical null distribution through iterative sampling from the data without assuming a known distribution, making it well-suited for unbalanced designs or non-parametric tests (Hayes, 1996). Given that the ISC data are in matrix form, we conducted a permutation test (Chen et al., 2016) by shuffling and resampling columns of the correlation matrix rather than using random permutations. This process was iterated for 100,000 iterations, and the threshold for the ISC-valence rating correlation coefficient was set at *p <* 0.000005 (α = 0.00001). This threshold corresponds to the two-tailed critical value within the distribution of ISC-valence rating correlation coefficients derived from resampled data.

### 2.4. Multidimensional scaling (MDS)

In this study, we employed multidimensional scaling (MDS) to explore whether the correlation coefficients between ISC and valence are linked to valence representation (Douglas Carroll & Arabie, 1998). MDS uses distance information to understand and visually represent the structure of high-dimensional data at a lower dimensional space. In psychological research, MDS is applied to evaluate whether objects within a set (such as stimuli, brain regions, and individuals) are appropriately positioned in the intended a low-dimensional space by computing distances between objects (Shinkareva et al., 2013). MDS results will demonstrate that objects with clear differences in characteristics are placed at greater distances from each other in this simplified space. On the other hand, objects sharing similar features are positioned closer together. This arrangement reflects how accurately the chosen features capture and represent the inherent attributes of the objects in consideration. We used MDS to confirm significant ROIs to effectively capture and explain the emotional dimensions related to valence.

We first extracted the dataset for the ROIs that corresponded with significant ISC-valence correlation coefficients (Figure 3 A). The voxel data related to the extracted ROI were formatted similarly to the matrix used for ISC calculation (621 segments by voxels) (Figure 3 B). The correlation matrix was calculated by using the voxel activity patterns for the 621 segments as displayed in Figure 3 C. Individual correlation matrices were computed and averaged across all 17 participants. The averaged matrix was then transformed into a distance matrix by subtracting it from a matrix consisting of all elements equal to 1 (]_621X621_) (Figure 3 D). Finally, the MDS solution was computed based on the distance matrix (Figure 3 E).

**Figure 3.**
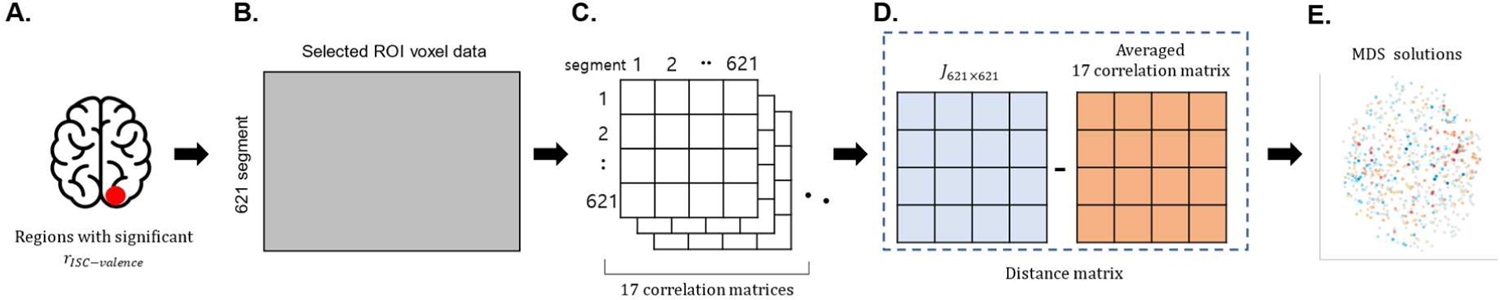
Illustration of the MDS calculation procedure

Procrustes rotation was used on MDS coordinates to establish valence as the primary dimension in this study. This rotation enabled the MDS values to align with the valence axis by using the participant’s average emotion ratings for each segment as the rotation criteria. Therefore, if the selected ROIs were implicated in emotion representation, the MDS solution for segments with positive content would be positioned to the right, while the MDS solution for segments with negative content would be positioned to the left. Moreover, the correlation coefficient between mean emotion ratings and the rotated MDS values was computed to statistically assess how well the rotated MDS coordinates represented the valence dimension. Furthermore, a permutation test was conducted to determine the significance of the correlation between MDS coordinates and valence ratings that were iterated 100,000 times. When we apply distance-based methods to convert high-dimensional data into lower dimensions, the resulting distribution deviates from the usual pattern seen in traditional linear models (Hajderanj et al., 2020). In this context, we established the threshold for the MDS-valence rating correlation coefficient at the top 5% (α = 0.05) to assess statistical significance.

### 2.6. Data visualization

In SPM12,3D Neuroimaging informatics technology initiative (NIfTI) files were generated to depict significant ISC-valence correlation coefficients. These files were subsequently employed in the MRIcroGL (v1.2.20220720) program, utilizing the MNI 152 template map for visualization.

The correlation coefficients between ISC and emotion ratings were displayed on both a 3D brain map and a 2D axial plane, employing mosaic displays as arbitrarily chosen with z coordinates of −10, 0, 10, 20, and 30. In regions exhibiting positive correlations, the coefficients were presented in a gradient ranging from dark red to light red, corresponding to an increase in correlation values (Figure 4 B). Conversely, for negative correlations, the coefficients were illustrated in a gradient from dark blue to light blue, reflecting a decrease in correlation values (Figure 4 C).

**Figure 4.**
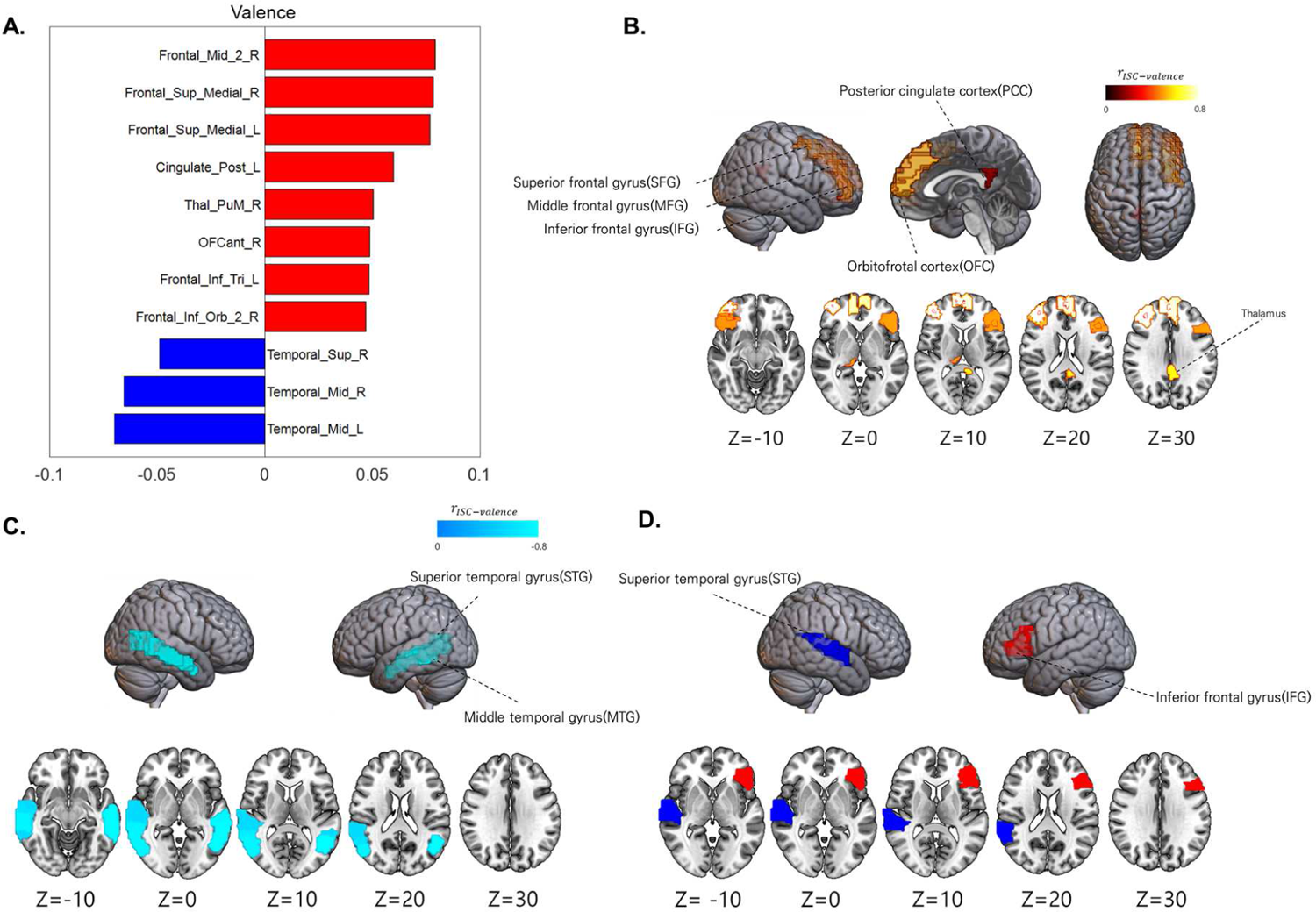
ISC-valence correlation coefficient results and MDS-valence correlation coefficient results. (A) Bar graphs depicting statistically significant ISC-valence correlations, (B) Regions indicating positive significance in ISC-valence correlations, (C) Regions indicating negative significance in ISC-valence correlations, and (D) Regions where both ISC-valence and MDS-valence correlations are statistically significant (red: positive, blue: negative).

Furthermore, the MDS-valence rating correlation coefficients were visualized on the MNI 152 template map using a similar approach as employed for ISC-valence correlation coefficients. By adjusting settings in MRIcroGL, areas where both ISC-valence and MDS-valence correlation coefficients showed statistically significant positive relationships were consistently highlighted in red. Similarly, regions where both ISC-valence and MDS-valence correlation coefficients were negatively significant were consistently visualized in blue (Figure 4 D).

## 3. Results

The results for the ISC-valence rating correlation coefficients revealed significant clusters including the right middle frontal gyrus (MFG) (r_ISC-valence_ = .079, *p <* .001), bilateral superior frontal gyrus medial (SFG) (central portion) r_ISC-valence_= .077, *p <* .001), left posterior cingulate cortex (PCC) (r = .06, *p <* .001), thalamus pulvinar medial (r_ISC-valence_ = .05, *p <* .001), right anterior orbital gyrus (r = .049, *p <* .001), and bilateral inferior frontal gyrus (IFG) (r_ISC-valence_ = .047, *p <* .001) regions (Figure 4 A, B). Furthermore, negative correlations were observed in the middle temporal gyrus (MTG) (r_ISC-valence_ = −.067, *p <* .001) and right hemisphere superior temporal gyrus (STG) (r_ISC-valence_ = −.049, *p <* .001) regions (Figure 4 A, C).

To examine the association between significant areas in ISC and actual valence representation, we examined the correlation between the rotated MDS values and the average emotion ratings for each segment. Permutation tests with 100,000 iterations were performed at a significance level of 0.05 using one-tailed tests. Among the regions where ISC-valence correlation coefficients showed significant positive relationships, the left IFG (r_MDS-valence_ = .047, *p <* .05) displayed a significant correlation with the MDS-valence rating coefficient. Similarly, within the regions where ISC-valence correlation coefficients were negative, the right STG ( r_MDS-valence_ = .092, *p <* .05) showed a significant correlation with the MDS coefficient (Figure 4 D).

## 4. Discussion

In this study, we investigated the consistency of neural responses among participants in specific regions related to valence. The significant correlation coefficients between ISC and valence ratings indicated heightened neural synchrony in response to emotional changes. Moreover, we employed MDS to validate significant regions.

In both the left IFG and right STG, significant correlations were observed between ISC-valence correlation coefficients and MDS-valence correlation coefficients. Previous research on humor-related studies has indicated that the IFG is linked to the comprehension and evaluation of jokes (Marinkovic et al., 2011), humor comprehension (Bartolo et al., 2006), understanding humor in cartoons (Campbell et al., 2015), and interpreting humor in social interactions (Uekermann et al., 2007). Grecucci et al. (2013) proposed that the IFG, in conjunction with the anterior insula and middle frontal gyrus, plays a crucial role in emotion reappraisal, interpreting others’ intentions, and and making decisions regarding emotional responses, contributing to interpersonal relationships and social interaction. In the stimuli, the character, Sherlock Holmes, applies his exceptional deductive abilities to reinterpret commonplace situations, deriving meaningful insights from the belongings and surroundings of his colleagues. Participants, influenced by Sherlock’s deductive inferences, may perceive ordinary scenes as a positively. Consequently, we speculated that participants perceived the scenes positively, leading to an increased neural synchrony in the IFG associated with reappraisal. This suggests that the IFG may play a crucial role in processing emotionally nuanced content, particularly when influenced by deductive reasoning cues.

The STG has previously been acknowledged for its role in multisensory integration (Beauchamp et al., 2004; Hagan et al., 2009) and integration of emotional information from various sensory modalities (Gao et al., 2019; Gao et al., 2020; Kim et al., 2016). While earlier studies concentrated on individual sensory cues, Lee Masson and Isik (2021) utilized fMRI data from two movie stimuli to explore brain regions associated with social interaction within a realistic visual environment. Their findings revealed that the STG exhibited distinctive responses to negative social situations, suggesting a specialized circuitry for engaging in negative interactions with others. The present study aligns with this prior research, as we observed robust neural synchrony in the STG when participants viewed emotionally charged, negative scenes. This suggests that participants may have interpreted conflictual interactions between characters in the “Sherlock” drama as real-world instances of negative social situations. Consistency with previous studies emphasizes the key role of the STG when figuring out how to handle emotional situations with other people, specifically in the context of negative social interactions.

Furthermore, the ISC analysis revealed heightened neural synchrony in the MFG, SFG, orbital frontal cortex (OFC), PCC, and IFG as the emotion shifted towards positivity. This outcome implies that participants perceived positive scenes in the video stimuli similarly, interpreting them as rewarding or favorable. This shared interpretation may have led to similar activation patterns in the SFG and OFC, which are known for their involvement in reward-related cognitive functions (Blair et al., 2007; Kringelbach, 2005; Kringelbach et al., 2003; Winecoff et al., 2013).

On the other hand, the activation pattern of the PCC may be explained by the characteristics of naturalistic stimuli. According to previous studies (Andreasen et al., 1995; Grasby et al., 1993), PCC is associated with memory, but another study (Maddock et al., 2003) indicates its activation is specific to emotional experiences rather than all types of memories. In our study, the stimuli were episodes from “Sherlock” where participants could recall memories or were be reminded of character backgrounds. This could have resulted in similar PCC activation patterns.

As the scenes became more negative, participants’ neural responses became more consistent in the bilateral MTG and the right STG. MTG is known to integrate cognitive and emotional information from stimuli, such as faces and voices. Goldin et al. (2008) conducted a study using short film clips inducing negative emotions for 15 seconds to investigate the neurobiology and regulation of emotion. They found that early (0-4.5 seconds) emotion processing involved the medial and left aspects of the PFC, while later emotion processing (10.5-15 seconds) decreased activity in the amygdala and PCC. They suggested that PFC underwent initial changes during the processing of valence that later influences the PFC, IFG and STG for emotional regulation. Therefore, the increased neural synchrony observed in the frontal regions in our study may be attributed to participants’ engagement in emotional reappraisal.

Our study identified neural synchrony in regions associated with emotional evaluation and social interaction, extending beyond the regions identified in previous ISC studies. A potential explanation for this difference lies in the characteristics of the stimuli presented. Nummenmaa et al. (2012) performed ISC while excluding auditory stimuli, providing movies without sound to native Finnish speakers. Participants were also given scripts explaining the content of the clips before watching. These experimental procedures might have hindered participants from fully understanding the cultural context of the stimuli or forming their own judgments about the situations. Consequently, we speculate that the naturalistic stimuli in Nummenmaa et al. (2012) may fail to fully capture the properties of stimuli in real-world scenarios, which may account for the absence of neural synchrony in regions related to memory, social interaction, and emotional reappraisal as observed in our present study.

### 4.1. Significance of the study

Our study explores the importance of observed neural synchrony in regions associated with valence, revealing insights into the complex processes involved in emotional responses and cognitive engagement. The increased consistency in neural responses, especially in the IFG and STG, indicates the significant role of these regions in processing emotional nuances that are influenced by deductive reasoning cues and negative social interactions.

The identification of neural synchrony in the MFG, SFG, OFC, PCC, and IFG during positive scenes emphasizes the shared perception of rewarding or favorable content among participants. This not only demonstrates the involvement of specific brain regions in reward-related cognitive functions but also provides valuable insights into the interplay between emotional processing and cognitive appraisal in positive contexts.

Moreover, our study furthers to the understanding of how naturalistic stimuli impact neural responses. The observed neural synchrony in regions related to memory, social interaction, and emotional reappraisal suggests that real-world scenarios, as portrayed in “Sherlock,” trigger intricate cognitive and emotional processes. This nuanced perspective deepens our comprehension of how individuals react to stimuli in a more realistic and culturally embedded context.

Previous experimental studies aimed to establish clear causal relationships between independent and dependent variables using controlled stimuli. In fMRI studies on emotions, various stimuli such as faces (Mourao-Miranda et al., 2011), pictures (Baucom et al., 2012), and sounds (Fruhholz et al., 2016) were utilized. However, they often deviated from real-life situations due to extreme manipulations. In our study, we examined how people’s neural responses changed in real-life situations in response to emotional changes by using a more realistic stimulus - the television drama “Sherlock.”

Furthermore, we addressed the challenges posed by the extensive duration of naturalistic stimuli in our study. Traditional fMRI data analysis methods, like the general linear model, face decreased performance or increased prediction uncertainty when applied to data measured over extended periods due to time-series autocorrelation. To overcome this, we applied new analysis methodologies such as ISC, ISC-valence rating correlation coefficients, and MDS to naturalistic stimuli, validating their effectiveness through permutation tests.

Our study utilized ISC to provide valuable complementary information on neural deactivation in fMRI data analysis. While it is commonly assumed that psychological changes induced by specific stimuli affect neural activation, interpreting decreased BOLD signals as decreased activation in brain regions associated with specific stimuli is challenging (Yuan et al., 2011). The ISC analysis method measures participants’ neural synchrony, capturing consistent patterns of neural activation or deactivation among participants, potentially contributing information not explored in other studies.

### 4.2. Limitations and suggestions

While our study benefits from utilizing naturalistic stimuli, which offer a closer representation of real-life scenarios, their inclusion introduces complexities in manipulating variables and establishing causal relationships. Unlike highly controlled stimuli, naturalistic stimuli lack precise control over emotional content, posing challenges in isolating specific factors of interest. For instance, a film clip portraying a dramatic event may elicit a multitude of emotions simultaneously, complicating the interpretation of emotional responses. Furthermore, the variability in contexts and content of naturalistic stimuli may result in individual differences in emotional experiences. Future research could address these issues through meta-analyses of emotion studies employing naturalistic stimuli, thus shedding light on the nuances of emotional processing in real-world settings.

In this study, correlation coefficients were computed for each region of interest, offering computational efficiency and compensating for equipment limitations. However, this approach analyzes fewer volumes compared to whole-brain data analysis. Although the AAL 3 template ensured accurate ROI anatomical localization, it may not have captured changes in neural synchrony outside the specific ROIs. The nearest-neighbor interpolation method used for ROI setting introduces discrepancies between actual anatomical locations and those in the study. To address this, future research could employ techniques such as searchlight analysis (Kriegeskorte et al., 2006) to continuously explore localized brain areas and compare voxel differences.

### 4.3. Future study

For future studies, it is imperative to give attention to both valence and arousal. An integral aspect of emotional research involves investigating the interaction between these core emotional dimensions. Notably, datasets of emotional stimuli, such as the International Affective Picture System (Lang et al., 1988) and the International Affective Digitized Sounds databases (Bradley & Lang, 2000), often exhibit uneven distributions, with positive and negative stimuli typically inducing higher levels of arousal compared to neutral stimuli. However, some of research on brain function suggests that distinct brain regions process valence and arousal separately (Barrett & Bliss-Moreau, 2009). For instance, the amygdala’s function is not to encode fear or negativity per se but rather to direct attention toward sensory stimuli of uncertain predictive value (Barad et al., 2006; Barrett et al., 2007; Holland & Gallagher, 1999; Wright et al., 2007). Similarly, the ventral striatum is responsible for coding reward or positivity through dopamine pathways (Berridge & Robinson, 1998; Horvitz, 2002; Salamone, 2009).

Initially, our aim was to identify brain regions sensitive to changes in valence induced by naturalistic stimuli. However, upon reviewing our study’s findings, we noticed significant involvement of brain regions where affective information integration occurs. We hypothesized that the processing of rich information inherent in naturalistic stimuli led to more intricate emotional processing, resulting in heightened neural activity in those areas. This underscores the complexity of emotional processing and highlights the necessity of considering interactions between valence and arousal dimensions within specific brain circuits when analyzing naturalistic stimuli.

While our study focused solely on valence, overlooking arousal, future research should integrate arousal data to attain a more comprehensive understanding of how these emotional dimensions interact in response to real-world stimuli. By examining both valence and arousal, we can better elucidate the nuanced mechanisms underlying emotional experiences and their neural substrates.

## 5. Data availability

This study is based on Sherlock Movie Watching Dataset (Chen et al., 2017). The data are available in follow link at https://dataspace.princeton.edu/handle/88435/dsp01nz8062179. These da ta were derived f rom the following resources available in the public domain: https://www.nature.com/articles/nn.4450, doi: https://doi.org/10.1038/nn.4450.

## <supplementary>

**Supplementary figure 1.**
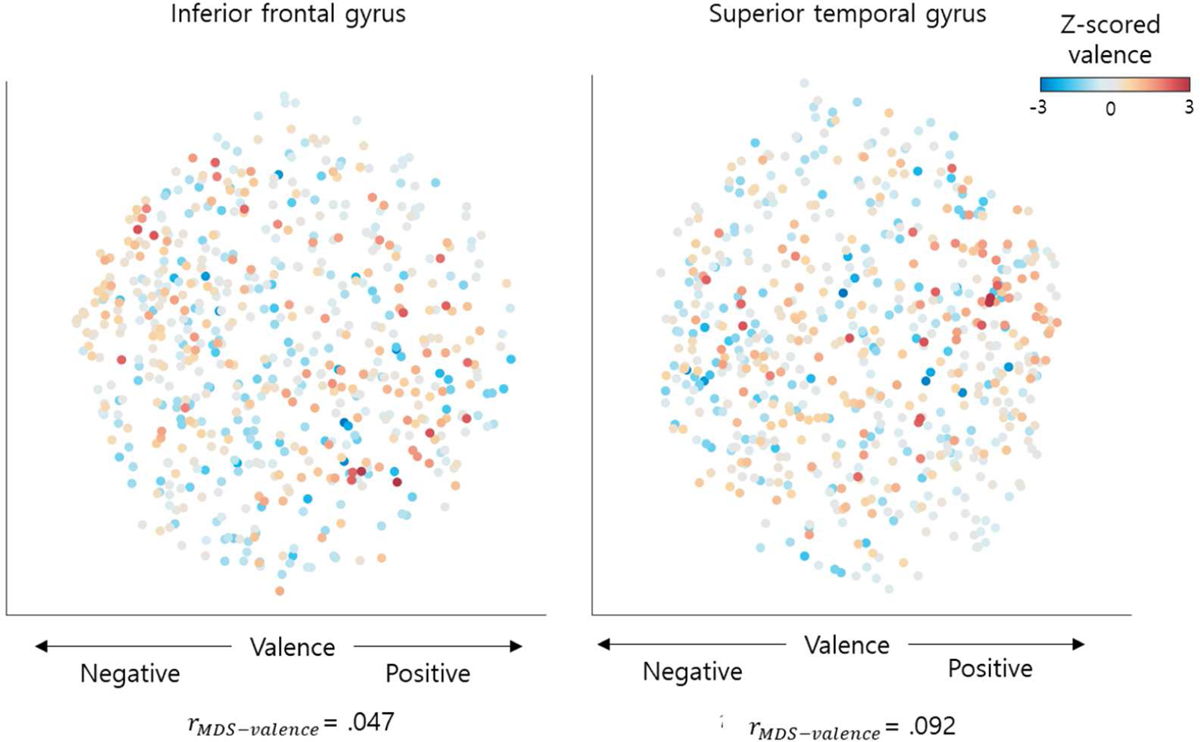
MDS visualization results. This figure shows the results obtained by MDS of the degree of neural activation corresponding to each segment. Each dot represents each segment, colored based on a z-score of valence ratings. In the coordinate plane, the x-axis reflects the valence dimension, and the y-axis represents a second dimension without rotation. The r value is the correlation coefficient of the mean valence ratings with the rotated MDS coordinates.

## Reference

Andreasen, N. C., O’Leary, D. S., Cizadlo, T., Arndt, S., Rezai, K., Watkins, G. L., Ponto, L. L., & Hichwa, R. D. (1995). Remembering the past: two facets of episodic memory explored with positron emission tomography. Am J Psychiatry, 152(11), 1576–1585. 10.1176/ajp.152.11.1576

Baldassano, C., Chen, J., Zadbood, A., Pillow, J. W., Hasson, U., & Norman, K. A. (2017). Discovering Event Structure in Continuous Narrative Perception and Memory. Neuron, 95(3), 709–721 e705. 10.1016/j.neuron.2017.06.041

Barad, M., Gean, P. W., & Lutz, B. (2006). The role of the amygdala in the extinction of conditioned fear. Biol Psychiatry, 60(4), 322–328. 10.1016/j.biopsych.2006.05.029

Barrett, L. F., & Bliss-Moreau, E. (2009). Affect as a Psychological Primitive. Adv Exp Soc Psychol, 41, 167–218. 10.1016/S0065-2601(08)00404-8

Barrett, L. F., Bliss-Moreau, E., Duncan, S. L., Rauch, S. L., & Wright, C. I. (2007). The amygdala and the experience of affect. Soc Cogn Affect Neurosci, 2(2), 73–83. 10.1093/scan/nsl042

Barrett, L. F., Quigley, K. S., Bliss-Moreau, E., & Aronson, K. R. (2004). Interoceptive sensitivity and self-reports of emotional experience. J Pers Soc Psychol, 87(5), 684–697. 10.1037/0022-3514.87.5.684

Bartolo, A., Benuzzi, F., Nocetti, L., Baraldi, P., & Nichelli, P. (2006). Humor comprehension and appreciation: an FMRI study. J Cogn Neurosci, 18(11), 1789–1798. 10.1162/jocn.2006.18.11.1789

Baucom, L. B., Wedell, D. H., Wang, J., Blitzer, D. N., & Shinkareva, S. V. (2012). Decoding the neural representation of affective states. NeuroImage, 59(1), 718–727. 10.1016/j.neuroimage.2011.07.037

Beauchamp, M. S., Argall, B. D., Bodurka, J., Duyn, J. H., & Martin, A. (2004). Unraveling multisensory integration: patchy organization within human STS multisensory cortex. Nat Neurosci, 7(11), 1190–1192. 10.1038/nn1333

Bennett, C. M., Wolford, G. L., & Miller, M. B. (2009). The principled control of false positives in neuroimaging. Soc Cogn Affect Neurosci, 4(4), 417–422. 10.1093/scan/nsp053

Berntson, G. G., Norman, G. J., Bechara, A., Bruss, J., Tranel, D., & Cacioppo, J. T. (2011). The insula and evaluative processes. Psychol Sci, 22(1), 80–86. 10.1177/0956797610391097

Berridge, K. C., & Robinson, T. E. (1998). What is the role of dopamine in reward: hedonic impact, reward learning, or incentive salience? Brain research reviews, 28(3), 309–369. 10.1016/S0165-0173(98)00019-8

Blair, K. S., Smith, B. W., Mitchell, D. G., Morton, J., Vythilingam, M., Pessoa, L., Fridberg, D., Zametkin, A., Sturman, D., Nelson, E. E., Drevets, W. C., Pine, D. S., Martin, A., & Blair, R. J. (2007). Modulation of emotion by cognition and cognition by emotion. NeuroImage, 35(1), 430–440. 10.1016/j.neuroimage.2006.11.048

Bradley, M. M., & Lang, P. J. (2000). Affective reactions to acoustic stimuli. Psychophysiology, 37(2), 204–215. https://www.ncbi.nlm.nih.gov/pubmed/10731770

Campbell, D. W., Wallace, M. G., Modirrousta, M., Polimeni, J. O., McKeen, N. A., & Reiss, J. P. (2015). The neural basis of humour comprehension and humour appreciation: The roles of the temporoparietal junction and superior frontal gyrus. Neuropsychologia, 79(Pt A), 10–20. 10.1016/j.neuropsychologia.2015.10.013

Chang, L. J., Jolly, E., Cheong, J. H., Rapuano, K. M., Greenstein, N., Chen, P. A., & Manning, J. R. (2021). Endogenous variation in ventromedial prefrontal cortex state dynamics during naturalistic viewing reflects affective experience. Sci Adv, 7(17), 7129–7152. 10.1126/sciadv.abf7129

Chen, G., Shin, Y. W., Taylor, P. A., Glen, D. R., Reynolds, R. C., Israel, R. B., & Cox, R. W. (2016). Untangling the relatedness among correlations, part I: Nonparametric approaches to inter-subject correlation analysis at the group level. NeuroImage, 142, 248–259. 10.1016/j.neuroimage.2016.05.023

Chen, J., Leong, Y. C., Honey, C. J., Yong, C. H., Norman, K. A., & Hasson, U. (2017). Shared memories reveal shared structure in neural activity across individuals. Nat Neurosci, 20(1), 115–125. 10.1038/nn.4450

Douglas Carroll, J., & Arabie, P. (1998). Multidimensional Scaling. In M. H. Birnbaum (Ed.), Measurement, judgment and decision making (pp. 179-250). Academic Press. 10.1016/b978-012099975-0.50005-1

Finn, E. S., Glerean, E., Khojandi, A. Y., Nielson, D., Molfese, P. J., Handwerker, D. A., & Bandettini, P. A. (2020). Idiosynchrony: From shared responses to individual differences during naturalistic neuroimaging. NeuroImage, 215, 116828. 10.1016/j.neuroimage.2020.116828

Friston, K. J., Holmes, A. P., Poline, J. B., Grasby, P. J., Williams, S. C., Frackowiak, R. S., & Turner, R. (1995). Analysis of fMRI time-series revisited. NeuroImage, 2(1), 45–53. 10.1006/nimg.1995.1007

Fruhholz, S., Trost, W., & Kotz, S. A. (2016). The sound of emotions-Towards a unifying neural network perspective of affective sound processing. Neurosci Biobehav Rev, 68, 96–110. 10.1016/j.neubiorev.2016.05.002

Gao, C., Weber, C. E., & Shinkareva, S. V. (2019). The brain basis of audiovisual affective processing: Evidence from a coordinate-based activation likelihood estimation meta-analysis. Cortex, 120, 66–77. 10.1016/j.cortex.2019.05.016

Gao, C., Weber, C. E., Wedell, D. H., & Shinkareva, S. V. (2020). An fMRI Study of Affective Congruence across Visual and Auditory Modalities. J Cogn Neurosci, 32(7), 1251–1262. 10.1162/jocn_a_01553

Goldin, P. R., McRae, K., Ramel, W., & Gross, J. J. (2008). The neural bases of emotion regulation: reappraisal and suppression of negative emotion. Biol Psychiatry, 63(6), 577–586. 10.1016/j.biopsych.2007.05.031

Grasby, P. M., Frith, C. D., Friston, K. J., Bench, C., Frackowiak, R. S., & Dolan, R. J. (1993). Functional mapping of brain areas implicated in auditory--verbal memory function. Brain, 116 (Pt 1)(1), 1-20. 10.1093/brain/116.1.1

Grecucci, A., Giorgetta, C., Bonini, N., & Sanfey, A. G. (2013). Reappraising social emotions: the role of inferior frontal gyrus, temporo-parietal junction and insula in interpersonal emotion regulation. Front Hum Neurosci, 7(SEP), 523. 10.3389/fnhum.2013.00523

Hagan, C. C., Woods, W., Johnson, S., Calder, A. J., Green, G. G., & Young, A. W. (2009). MEG demonstrates a supra-additive response to facial and vocal emotion in the right superior temporal sulcus. Proc Natl Acad Sci U S A, 106(47), 20010–20015. 10.1073/pnas.0905792106

Hajderanj, L., Chen, D. Q., Grisan, E., & Dudley, S. (2020). Single- and Multi-Distribution Dimensionality Reduction Approaches for a Better Data Structure Capturing. IEEE Access, 8, 207141–207155. 10.1109/Access.2020.3038460

Hasson, U., Malach, R., & Heeger, D. J. (2010). Reliability of cortical activity during natural stimulation. Trends Cogn Sci, 14(1), 40–48. 10.1016/j.tics.2009.10.011

Hasson, U., Nir, Y., Levy, I., Fuhrmann, G., & Malach, R. (2004). Intersubject synchronization of cortical activity during natural vision. Science, 303(5664), 1634–1640. 10.1126/science.1089506

Hasson, U., Yang, E., Vallines, I., Heeger, D. J., & Rubin, N. (2008). A hierarchy of temporal receptive windows in human cortex. J Neurosci, 28(10), 2539–2550. 10.1523/JNEUROSCI.5487-07.2008

Hayes, A. F. (1996). Permutation test is not distribution-free: Testing H₀: ρ = 0. Psychological Methods, 1(2), 184–198. 10.1037/1082-989x.1.2.184

Hejnar, M. P., Kiehl, K. A., & Calhoun, V. D. (2007). Interparticipant correlations: a model free FMRI analysis technique. Hum Brain Mapp, 28(9), 860–867. 10.1002/hbm.20321

Holland, P. C., & Gallagher, M. (1999). Amygdala circuitry in attentional and representational processes. Trends Cogn Sci, 3(2), 65–73. 10.1016/s1364-6613(98)01271-6

Honey, C. J., Thesen, T., Donner, T. H., Silbert, L. J., Carlson, C. E., Devinsky, O., Doyle, W. K., Rubin, N., Heeger, D. J., & Hasson, U. (2012). Slow cortical dynamics and the accumulation of information over long timescales. Neuron, 76(2), 423–434. 10.1016/j.neuron.2012.08.011

Horvitz, J. C. (2002). Dopamine, Parkinson’s disease, and volition. Behavioral and Brain Sciences, 25(5), 586–586. 10.1017/S0140525X02300104

Kauppi, J. P., Jaaskelainen, I. P., Sams, M., & Tohka, J. (2010). Inter-subject correlation of brain hemodynamic responses during watching a movie: localization in space and frequency. Front Neuroinform, 4(MAR), 5. 10.3389/fninf.2010.00005

Kayyal, H., Yiannakas, A., Kolatt Chandran, S., Khamaisy, M., Sharma, V., & Rosenblum, K. (2019). Activity of Insula to Basolateral Amygdala Projecting Neurons is Necessary and Sufficient for Taste Valence Representation. J Neurosci, 39(47), 9369–9382. 10.1523/JNEUROSCI.0752-19.2019

Kim, J., Shinkareva, S. V., & Wedell, D. H. (2017). Representations of modality-general valence for videos and music derived from fMRI data. NeuroImage, 148, 42–54. 10.1016/j.neuroimage.2017.01.002

Kim, J., Wang, J., Wedell, D. H., & Shinkareva, S. V. (2016). Identifying Core Affect in Individuals from fMRI Responses to Dynamic Naturalistic Audiovisual Stimuli. PLOS ONE, 11(9), e0161589. 10.1371/journal.pone.0161589

Kim, J., Weber, C. E., Gao, C., Schulteis, S., Wedell, D. H., & Shinkareva, S. V. (2020). A study in affect: Predicting valence from fMRI data. Neuropsychologia, 143, 107473. 10.1016/j.neuropsychologia.2020.107473

Kriegeskorte, N., Goebel, R., & Bandettini, P. (2006). Information-based functional brain mapping. Proc Natl Acad Sci U S A, 103(10), 3863–3868. 10.1073/pnas.0600244103

Kringelbach, M. L. (2005). The human orbitofrontal cortex: linking reward to hedonic experience. Nat Rev Neurosci, 6(9), 691–702. 10.1038/nrn1747

Kringelbach, M. L., O’Doherty, J., Rolls, E. T., & Andrews, C. (2003). Activation of the human orbitofrontal cortex to a liquid food stimulus is correlated with its subjective pleasantness. Cereb Cortex, 13(10), 1064–1071. 10.1093/cercor/13.10.1064

Lang, P. J., Bradley, M. M., & Cuthbert, B. N. (1988). International affective picture system. 10.1037/t66667-000

Lee Masson, H., & Isik, L. (2021). Functional selectivity for social interaction perception in the human superior temporal sulcus during natural viewing. NeuroImage, 245, 118741. 10.1016/j.neuroimage.2021.118741

Lerner, Y., Honey, C. J., Silbert, L. J., & Hasson, U. (2011). Topographic mapping of a hierarchy of temporal receptive windows using a narrated story. J Neurosci, 31(8), 2906–2915. 10.1523/JNEUROSCI.3684-10.2011

Li, X., Zhu, Y., Vuoriainen, E., Ye, C., & Astikainen, P. (2021). Decreased intersubject synchrony in dynamic valence ratings of sad movie contents in dysphoric individuals. Sci Rep, 11(1), 14419. 10.1038/s41598-021-93825-1

Maddock, R. J., Garrett, A. S., & Buonocore, M. H. (2003). Posterior cingulate cortex activation by emotional words: fMRI evidence from a valence decision task. Hum Brain Mapp, 18(1), 30–41. 10.1002/hbm.10075

Marinkovic, K., Baldwin, S., Courtney, M. G., Witzel, T., Dale, A. M., & Halgren, E. (2011). Right hemisphere has the last laugh: neural dynamics of joke appreciation. Cogn Affect Behav Neurosci, 11(1), 113–130. 10.3758/s13415-010-0017-7

Mesulam, M. M. (2000). Behavioral Neuroanatomy: Large-scale networks, association cortex, frontal syndromes, the limbic system, and hemispheric specializations. In Principles of behavioral and cognitive neurology, 2nd ed. (pp. 1-120). Oxford University Press.

Miskovic, V., & Anderson, A. K. (2018). Modality general and modality specific coding of hedonic valence. Curr Opin Behav Sci, 19, 91–97. 10.1016/j.cobeha.2017.12.012

Miyake, A., Friedman, N. P., Emerson, M. J., Witzki, A. H., Howerter, A., & Wager, T. D. (2000). The unity and diversity of executive functions and their contributions to complex “Frontal Lobe” tasks: a latent variable analysis. Cogn Psychol, 41(1), 49–100. 10.1006/cogp.1999.0734

Mourao-Miranda, J., Hardoon, D. R., Hahn, T., Marquand, A. F., Williams, S. C., Shawe-Taylor, J., & Brammer, M. (2011). Patient classification as an outlier detection problem: an application of the One-Class Support Vector Machine. NeuroImage, 58(3), 793–804. 10.1016/j.neuroimage.2011.06.042

Najafi, M., Kinnison, J., & Pessoa, L. (2017). Dynamics of Intersubject Brain Networks during Anxious Anticipation. Front Hum Neurosci, 11, 552. 10.3389/fnhum.2017.00552

Nastase, S. A., Gazzola, V., Hasson, U., & Keysers, C. (2019). Measuring shared responses across subjects using intersubject correlation. Soc Cogn Affect Neurosci, 14(6), 667–685. 10.1093/scan/nsz037

Nichols, T., & Hayasaka, S. (2003). Controlling the familywise error rate in functional neuroimaging: a comparative review. Stat Methods Med Res, 12(5), 419–446. 10.1191/0962280203sm341ra

Nichols, T. E. (2012). Multiple testing corrections, nonparametric methods, and random field theory. NeuroImage, 62(2), 811–815. 10.1016/j.neuroimage.2012.04.014

Nichols, T. E., & Holmes, A. P. (2002). Nonparametric permutation tests for functional neuroimaging: a primer with examples. Hum Brain Mapp, 15(1), 1–25. 10.1002/hbm.1058

Nummenmaa, L., Glerean, E., Viinikainen, M., Jaaskelainen, I. P., Hari, R., & Sams, M. (2012). Emotions promote social interaction by synchronizing brain activity across individuals. Proc Natl Acad Sci U S A, 109(24), 9599–9604. 10.1073/pnas.1206095109

Rolls, E. T., Huang, C. C., Lin, C. P., Feng, J., & Joliot, M. (2020). Automated anatomical labelling atlas 3. NeuroImage, 206, 116189. 10.1016/j.neuroimage.2019.116189

Russell, J. A., Weiss, A., & Mendelsohn, G. A. (1989). Affect Grid - a Single-Item Scale of Pleasure and Arousal. Journal of Personality and Social Psychology, 57(3), 493–502. 10.1037/0022-3514.57.3.493

Salamone, J. D. (2009). Dopamine, effort, and decision making: theoretical comment on Bardgett et al. (2009). Behav Neurosci, 123(2), 463-467. 10.1037/a0015381

Schmuckler, M. A. (2001). What Is Ecological Validity? A Dimensional Analysis. Infancy, 2(4), 419–436. 10.1207/S15327078IN0204_02

Shin, L. M., Wright, C. I., Cannistraro, P. A., Wedig, M. M., McMullin, K., Martis, B., Macklin, M. L., Lasko, N. B., Cavanagh, S. R., Krangel, T. S., Orr, S. P., Pitman, R. K., Whalen, P. J., & Rauch, S. L. (2005). A functional magnetic resonance imaging study of amygdala and medial prefrontal cortex responses to overtly presented fearful faces in posttraumatic stress disorder. Arch Gen Psychiatry, 62(3), 273–281. 10.1001/archpsyc.62.3.273

Shinkareva, S. V., Wang, J., & Wedell, D. H. (2013). Examining similarity structure: multidimensional scaling and related approaches in neuroimaging. Comput Math Methods Med, 2013, 796183. 10.1155/2013/796183

Simony, E., Honey, C. J., Chen, J., Lositsky, O., Yeshurun, Y., Wiesel, A., & Hasson, U. (2016). Dynamic reconfiguration of the default mode network during narrative comprehension. Nat Commun, 7, 12141. 10.1038/ncomms12141

Sonkusare, S., Breakspear, M., & Guo, C. (2019). Naturalistic Stimuli in Neuroscience: Critically Acclaimed. Trends Cogn Sci, 23(8), 699–714. 10.1016/j.tics.2019.05.004

Turner, B. M., Forstmann, B. U., Love, B. C., Palmeri, T. J., & Van Maanen, L. (2017). Approaches to Analysis in Model-based Cognitive Neuroscience. J Math Psychol, 76(B), 65–79. 10.1016/j.jmp.2016.01.001

Uekermann, J., Daum, I., & Channon, S. (2007). Toward a cognitive and social neuroscience of humor processing. Social Cognition, 25(4), 553–572. 10.1521/soco.2007.25.4.553

van Baar, J. M., Chang, L. J., & Sanfey, A. G. (2019). The computational and neural substrates of moral strategies in social decision-making. Nat Commun, 10(1), 1483 10.1038/s41467-019-09161-6

Vicente, R., Wibral, M., Lindner, M., & Pipa, G. (2011). Transfer entropy--a model-free measure of effective connectivity for the neurosciences. J Comput Neurosci, 30(1), 45–67. 10.1007/s10827-010-0262-3

Wang, J., Conder, J. A., Blitzer, D. N., & Shinkareva, S. V. (2010). Neural representation of abstract and concrete concepts: a meta-analysis of neuroimaging studies. Hum Brain Mapp, 31(10), 1459–1468. 10.1002/hbm.20950

Winecoff, A., Clithero, J. A., Carter, R. M., Bergman, S. R., Wang, L., & Huettel, S. A. (2013). Ventromedial prefrontal cortex encodes emotional value. J Neurosci, 33(27), 11032–11039. 10.1523/JNEUROSCI.4317-12.2013

Wright, C. I., Dickerson, B. C., Feczko, E., Negeira, A., & Williams, D. (2007). A functional magnetic resonance imaging study of amygdala responses to human faces in aging and mild Alzheimer’s disease. Biol Psychiatry, 62(12), 1388–1395. 10.1016/j.biopsych.2006.11.013

Yuan, H., Perdoni, C., Yang, L., & He, B. (2011). Differential electrophysiological coupling for positive and negative BOLD responses during unilateral hand movements. J Neurosci, 31(26), 9585–9593. 10.1523/JNEUROSCI.5312-10.2011

